# A novel predicted ADP-ribosyltransferase family conserved in eukaryotic evolution

**DOI:** 10.1101/2020.07.21.169896

**Authors:** Zbigniew Wyżewski, Marcin Gradowski, Marianna Krysińska, Małgorzata Dudkiewicz, Krzysztof Pawłowski

## Abstract

The presence of many completely uncharacterized proteins, even in well-studied organisms such as humans, seriously hampers full understanding of the functioning of the living cells. ADP-ribosylation is a common post-translational modification of proteins; also nucleic acids and small molecules can be modified by the covalent attachment of ADP-ribose. This modification, important in cellular signalling and infection processes, is usually executed by enzymes from the large superfamily of ADP-ribosyltransferases (ARTs)

Here, using bioinformatics approaches, we identify a novel putative ADP-ribosyltransferase family, conserved in eukaryotic evolution, with a divergent active site. The hallmark of these proteins is the ART domain nestled between flanking leucine-rich repeat (LRR) domains. LRRs are involved in innate immune surveillance.

The novel family appears as likely novel ADP-ribosylation “writers”, previously unnoticed new players in cell signaling by this emerging post-translational modification. We propose that this family, including its human member LRRC9, may be involved in an ancient defense mechanism, with analogies to the innate immune system, and coupling pathogen detection to ADP-ribosyltransfer signalling.

## Introduction

ADP-ribosyltransferases (ARTs) are enzymes catalyzing the transfer of ADP-ribose from oxidized nicotinamide adenine dinucleotide (NAD^+^) to different acceptor molecules. Thus, they are responsible for chemical modification of various target molecules such as proteins, nucleic acids and small molecules. The enzymes are highly conserved in evolution and widespread in nature. They are common to all three domains of life: the Archaea, the Bacteria and the Eukarya. Multiplicity of ADP-ribosylated substrates is reflected in a variety of functions performed by ARTs [1–4].

Prokaryotic ADP-ribosyltransferases often play a role of bacterial toxins or effectors. Several studies show that ADP-ribosylation of the host target molecules by ARTs from taxonomically diverse bacteria may be correlated with the development of infection as its essential step [5–9]. At the intracellular scale, it may change apoptotic potential of infected cell [10] and/or disturb organization of cellular membranes and actin cytoskeleton [11–13] whereas at systemic level, it can disturb cell-mediated immune response [6, 10], or increase the permeability of barriers limiting bacterial spread (epithelium and endothelium) [8, 10] and consequently contribute to serious structural and functional disorders of the host tissues and organs [14, 15]. Some ARTs act on small molecules, e.g. the bacterial Arr enzymes that ADP-ribosylate the antibiotics rifamycins [16].

Eukaryotic ADP-ribosyltransferases play important roles in both physiological and pathophysiological processes, contributing to change in chemical properties of the ADP-ribose acceptors. Proteins, post-translationally modified by ARTs, can also lose the capacity to interact with their ligands or acquire the ability to bind new ones. ADP-ribosylation of enzymes may cause substantial changes in their catalytic activity [17, 18]. ADP-ribose moieties may function as signals that direct modified acceptors to ubiquitin-dependent proteolysis [1, 3]. Influencing the half-life of the target molecules, ADP-ribosyltransferases determine their intracellular level and thus affect their activity [3]. Moreover, ADP-ribose moieties may form molecular scaffolds with a negative charge. Such structures play a role in recruitment of positively charged proteins, favoring specific intermolecular interactions [19]. Poly-ADP-ribosylation of DNA-binding proteins (e.g. core histones [20, 21]), chromatin remodeling enzymes (e.g. histone demethylase KDM5B [22], DEK protein [23] and DNA repair factors [24]) influences genome organization [21] and expression [22, 23] as well as effective DNA repair [20, 25]. Recently, growing evidence shows that PARPs also directly ADP-ribosylate mRNA [26]. Other examples of nucleic acid ADP-ribosylation are provided by the bacterial toxin DarT that acts on ssDNA [27] and the toxin scabin, acting on mononucleosides, nucleotides, and both single-stranded and dsDNA [28].

The members of the poly-ADP-ribosyltransferase (PARP) family, PARP1 and PARP2, are examples of ARTs that modify proteins interacting with DNA. PARP1 and PARP2 are localized in the nucleus where they are involved in many cellular processes [29–32]. PARP1 and PARP2 transferase activity is known to be necessary for repair of DNA damage such as deamination, hydroxylation and methylation of nitrogenous bases as well as for repair of single strand breaks. PARP1 and PARP also play a role in maintaining telomere stability[33, 34].

In general, ADP-ribosyltransferases are responsible for regulation of intracellular and extracellular signal transduction. Therefore, their activity determines viability and proliferation potential of cells, DNA stability, immune system reactivity and thus the proper functioning of eukaryotic organisms [33–36]. On the other hand, ADP-ribosyltransferases may also be involved in development of pathological states such as neurodegenerative disorders, diabetes, atherosclerosis, cataract and cancer [37–42]. Similarly to protein phosphorylation, ADP-ribosylation is a reversible post-translational modification, performed by a trio of “writers” (ARTs), “readers” (e.g. macro domain proteins) and “erasers” (e.g. some macro domains, ADP-ribose hydrolases, and NUDIX phosphodiesterases) [43].

According to the Pfam database, the ART clan (superfamily) comprises 19 families of domains, eight of which, ART, DUF952, Enterotoxin_a, PARP, Pertussis_S1, PTS_2-RNA, RolB_RolC and TNT, include eukaryotic members [44]. PARPs, the best studied family of ADP-ribosyltransferases, are responsible for modification of target structures by covalently adding polymeric chains of ADP-ribose moieties instead of transferring only one moiety. Despite low conservation of amino acid sequence, ART clan members are characterized by a common spatial structure comprising a split β-sheet and two helical regions surrounding it. The “split” separates the β-sheet into two halves, each composed of three β-strands (4-5-2 and 1-3-6, respectively). [1, 3]. Aravind and colleagues divide ARTs into three main clades: the H-H-h clade, the H-Y-[EDQ] clade and the R-S-E clade [45] that are characterized by the different configurations of active site amino acid residues. In the H-H-h clade domains, the catalytic centre comprises two histidines and one hydrophobic residue supplied by β-strands 1, 2 and 5, respectively. The H-Y-[EDQ] clade domains, including PARPs, are characterized by active site composed of histidine, tyrosine and glutamate/aspartate/glutamine whereas in the R-[ST]-E clade domains, active site triad comprises arginine, a polar residue (serine/threonine) and glutamate [1]. Three families of the ART clan, PARP, PTS_2-RNA and ART, are present in humans, and contain 16, 1 and 4 representatives, respectively. The NEURL-4 ART-like domain family [46], not included in Pfam database, is the fourth human ART-like family. The full catalogue of ADP-ribosylation enzymes is likely far from completion, as one may expect from recent discoveries of novel ART domains in effectors from pathogenic bacteria that perform non-canonical ubiquitination [5, 47, 48], novel ART/macro pairs in bacterial toxin/antitoxin systems [49] and novel human macro domains [50]. Other examples of the novel ADP-ribosylation players are provided by the viral macro domains, present in many dangerous viruses, including the SARS and SARS-CoV-2 coronaviruses [51], and by the recently characterized novel ART-like domain (DUF3715) in the human TASOR protein involved in gene silencing [52, 53].

In this paper, building up on experience in identification of novel enzyme families [50, 54–57], we identified structural similarity between ART-like catalytic domains and a family of eukaryotic uncharacterized protein domains present in homologs of the leucine-rich repeat containing protein 9 (LRRC9). For clarity, we will call this domain LRRC9-ART. We investigated sequence similarities between the novel and the known ADP-ribosyltransferases. We reconstructed phylogenetic relationships between sequences within LRRC9 domain family and other ART clan families. We also explored sequence variability in the novel ART-like domain in the context of predicted three-dimensional structures and proposed their likely biological functions.

## Materials and methods

The FFAS03 [58], HHpred [59] and Phyre2 [60] servers were used to determine distant sequence similarities of LRRC9 central domain to proteins of known structures from public databases. Standard parameters and significance thresholds were selected.

The representative set of LRRC9 ART-like domain sequences was collected by submitting the ART-like domain of the human LRRC9 protein (UniProtKB: Q6ZRR7.2, positions 389-628) to two iterations of jackhmmer (also with standard parameters) ran on the Reference Proteomes database.

The MAFFT program [61] was used to build the multiple sequence alignments of the novel domains and PARP catalytic domains obtained from the rp75 set from the Pfam database [44]. In-house scripts were used to merge family-wise multiple sequence alignments according to a FFAS pairwise alignment of representatives of the two families. The WebLogo server [62] was used to visualize results as sequence logos.

The CLANS [63] algorithm was used to visualize close and distant similarities between the putative and the known ADP-ribosyltransferase domains. Sequence similarities up to BLAST E-value of 1, 1e^−2^ or 1e^−4^ and the BLOSUM45 substitution matrix were used. In order to acquire collection of the known ADP-ribosyltransferase domains, the following sets of sequences were obtained from Pfam database: ADPrib_exo_Tox rp15, ADPRTs_Tse2 rp15, Anthrax-tox_M rp75, Arr-ms rp15, ART rp15, ART-PolyVal rp15, AvrPphF-ORF-2 rp15, Diphtheria_C rp75, Dot_icm_IcmQ rp35, DUF2441 rp15, DUF952 rp15, Enterotoxin_a rp15, Exotox-A_cataly rp75, NADase_NGA rp35, PARP rp15, Pertussis_S1 rp15, PTS_2-RNA rp15, RolB_RolC rp55 and TNT rp15. Next, collected data were supplemented with four sequence sets representing the following ART-like domain families not included in Pfam database: ESPJ, SidE, NEURL4 and DUF3715/TASOR. Sets of ESPJ and SidE domain sequences were obtained by submitting sequences from UniProtKB and Protein NCBI databases (UniProtKB: W0AL45, positions 52-214 and refseq ID: YP_094288.1, positions 686-912, respectively) to two iterations of jackhmmer with standard parameters, ran on UniProtKB database. The input sequence of NEURL4 (UniProtKB: F2TYZ7, positions 102-275) was subject to the same procedure, but extended to three iterations and ran on the Reference Proteomes database. Set of DUF3715 sequences was obtained by submitting the TASOR ART domain (UniProtKB: Q9UK61.3, positions 92-337) to two iterations of jackhmmer with standard parameters. Next, the whole collection of putative and known domain sequences were prepared for the CLANS procedure by clustering with CD-HIT and selecting representatives at a 70% sequence identity threshold [64].

In order to determine phylogenetic spread of the novel ART domains, a set of LRRC9 ART-like domain homologs was obtained using jackhmmer search seeded with input sequence UniProtKB: Q6ZRR7.2, positions 389-628. Then, Phylogeny.fr platform [65] was utilized to construct phylogenetic tree. A multiple sequence alignment was generated using the MUSCLE program [66]. Next, advanced mode of the platform was utilized to build phylogenetic tree using the maximum likelihood PhyML program [67] with standard settings (WAG amino-acid substitution model, four substitution rate categories, aLRT test for bootstrap support). Phylogenetic relationships between homologs were visualized as a dendrogram using the iTOL server [68].

Protein three-dimensional models for hypothetical structures of the human LRRC9-ART domain were built using homology modelling method Modeller9v21 [69] and were based on human PARP10 structure template (PDB ID: 3HKV).

Putative ligand binding site in the modelled structure was identified using the COFACTOR algorithm [70], an option delivered by the I-Tasser (Iterative Threading ASSEmbly Refinement) server for protein structure and function prediction [71] based on threading the query structural model through the BioLiP protein function database. COFACTOR identifies potential functional sites by local and global structure matches and suggests annotations. Structures were visualized and analyzed using UCSF Chimera tools [72].

The three-domain model for whole human LRRC9 sequence was built using AIDA (Ab Initio Domain Assembly Server) [45] based on structures modelled using Modeller9v21 separately for three identified domains: N-terminal LRR region with helical fragment (PDB code of modelling template: 3OJA), ART-like domain (template: 3HKV), and C-terminal LRR region domain (template: 4LSX).

Conservation values of sequence positions in multiple sequence alignment created for the human LRRC9-ART domain homologues (for mapping onto the modelled structures), were derived from Jalview alignment editor [73] and were automatically calculated as the number of conserved physicochemical properties for each column of the alignment [74].

The LRR repeats were identified using the LRRFinder tool [75]. Gene Ontology analysis of human genes encoding LRR-containing proteins (i.e. their cellular/extracellular localization, molecular function and association with biological processes) was performed with the use of Panther Classification System [76]. The domain organization of LRRC9 proteins was visualised using the DOG tool [77].

For analysis of protein and mRNA expression, the Proteomics-db and Protein Atlas databases were used [78, 79]. The BioGRID Interaction Database was used to obtain information about possible place of LRRC9 in human protein-protein interaction networks [80]. For prediction of the subcellular localization of LRRC9, the DeepLoc server was used [81].

## Results and discussion

### Assignment of the central domain of LRRC9 to the ADP-ribosyl clan

A sequence screen of human proteins for remote homologs of ADP-ribosyltransferases (ARTs) using the FFAS03 method indicated that weak ART similarity might exist in the central region of the LRRC9 protein (Fig. 1). In order to examine in detail this similarity, we used FFAS03, HHpred and Phyre2 servers. All of them are dedicated to protein structure prediction, allowing detection of non-obvious sequence similarity between very distant protein homologs. These three independent bioinformatics tools revealed statistically significant albeit distant sequence similarity of a region of human LRRC9 protein to representatives of the PARP catalytic domain family (Table 1). Such remote sequence similarity points to possible enzymatic function of the putative ART domain.

**Fig. 1.**
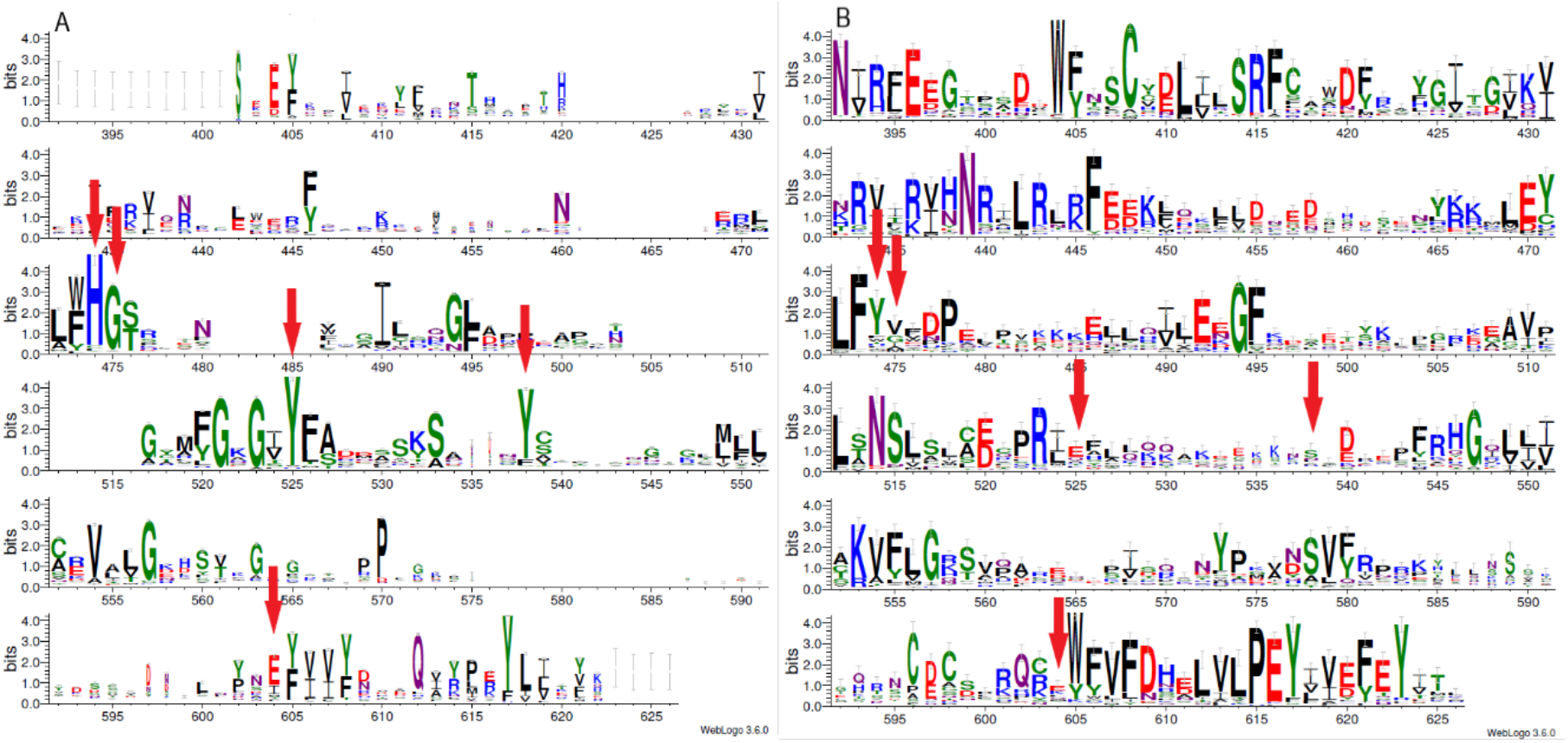
Sequence logos for PARP homologues (A) and LRRC9 homologues (B), created from MAFFT alignment using WebLogo server. Sequence numbering according to human LRRC9 protein. The two logos are aligned using pairwise FFAS03 alignment. The catalytically important PARP residues His, Gly, Tyr, Tyr and Glu (aligned to positions 474, 475, 525, 538 and 604, respectively, of the human LRRC9 protein) marked with red arrows.

**Table 1.**
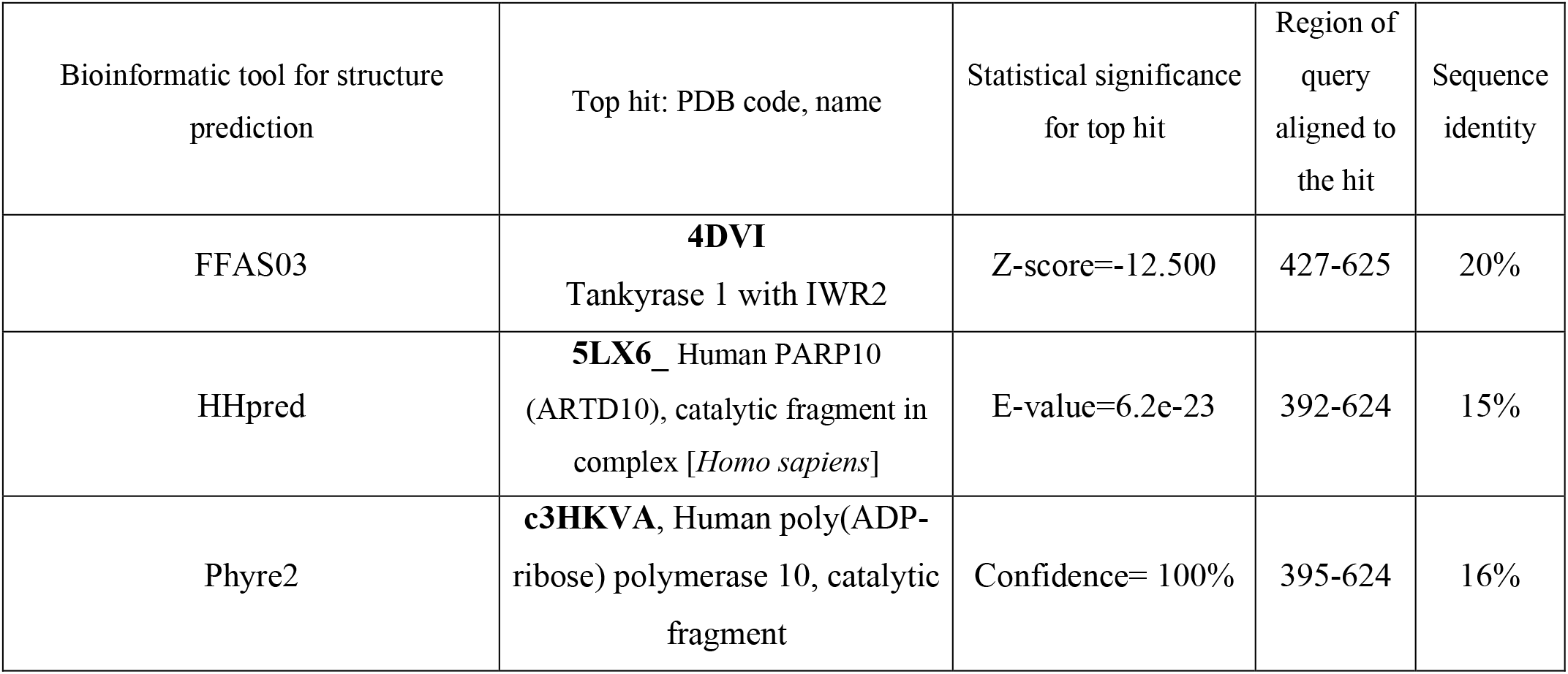
Top structure predictions for ART-like domain of the human LRRC9 protein

| Bioinformatic tool for structure prediction | Top hit: PDB code, name | Statistical significance for top hit | Region of query aligned to the hit | Sequence identity |
| --- | --- | --- | --- | --- |
| FFAS03 | **4DVI**<br>Tankyrase 1 with IWR2 | Z-score=-12.500 | 427-625 | 20% |
| HHpred | **5LX6_** Human PARP10<br>(ARTD10), catalytic fragment in complex [*Homo sapiens*] | E-value=6.2e-23 | 392-624 | 15% |
| Phyre2 | **c3HKVA**, Human poly(ADP-ribose) polymerase 10, catalytic fragment | Confidence= 100% | 395-624 | 16% |

The relationship of the novel LRRC9 ART-like family, found in many eukaryotic lineages (Fig. 2), to the known ART families can be visualized using the sequence-based CLANS clustering approach (Fig. 3). CLANS analysis was performed at three different E-value levels for BLAST hits for all known families containing ADP-ribosyltransferase domain. In agreement with the FFAS03, HHpred and Phyre results, CLANS results suggest a closer relation between the novel ART family and the PARP domains, noticeable at E-value 1, 1e^−2^ and even at 1e^−4^. This, together with sequence conservation analysis allowed us to hypothesize that LRRC9-ART belongs to the H-Y-[EDQ] clade.

**Fig. 2.**
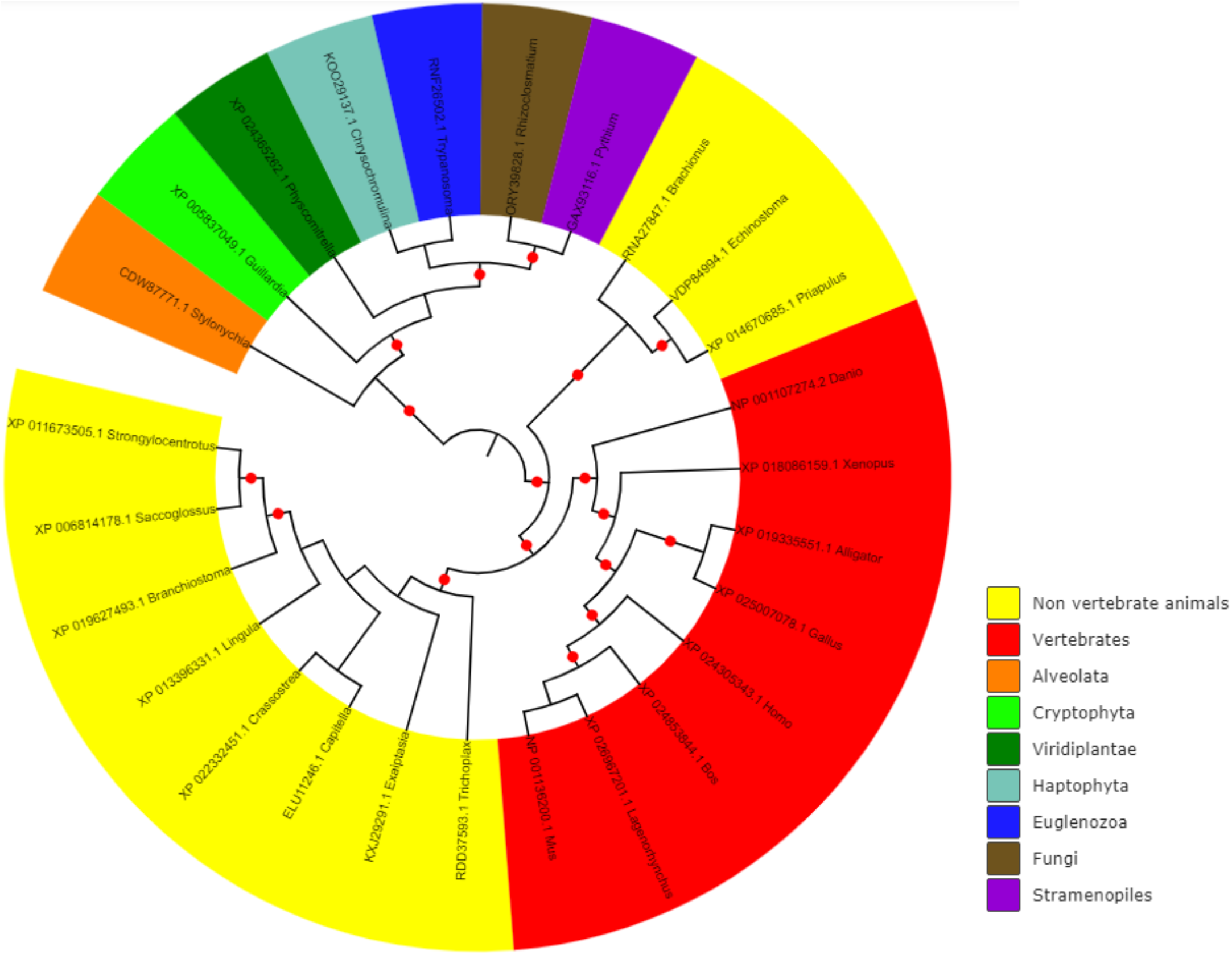
Phylogenetic spread of LRRC9 ART-like domain family. Maximum likelihood tree. Red circles mark bootstrap support of at least 75%.

**Fig. 3.**
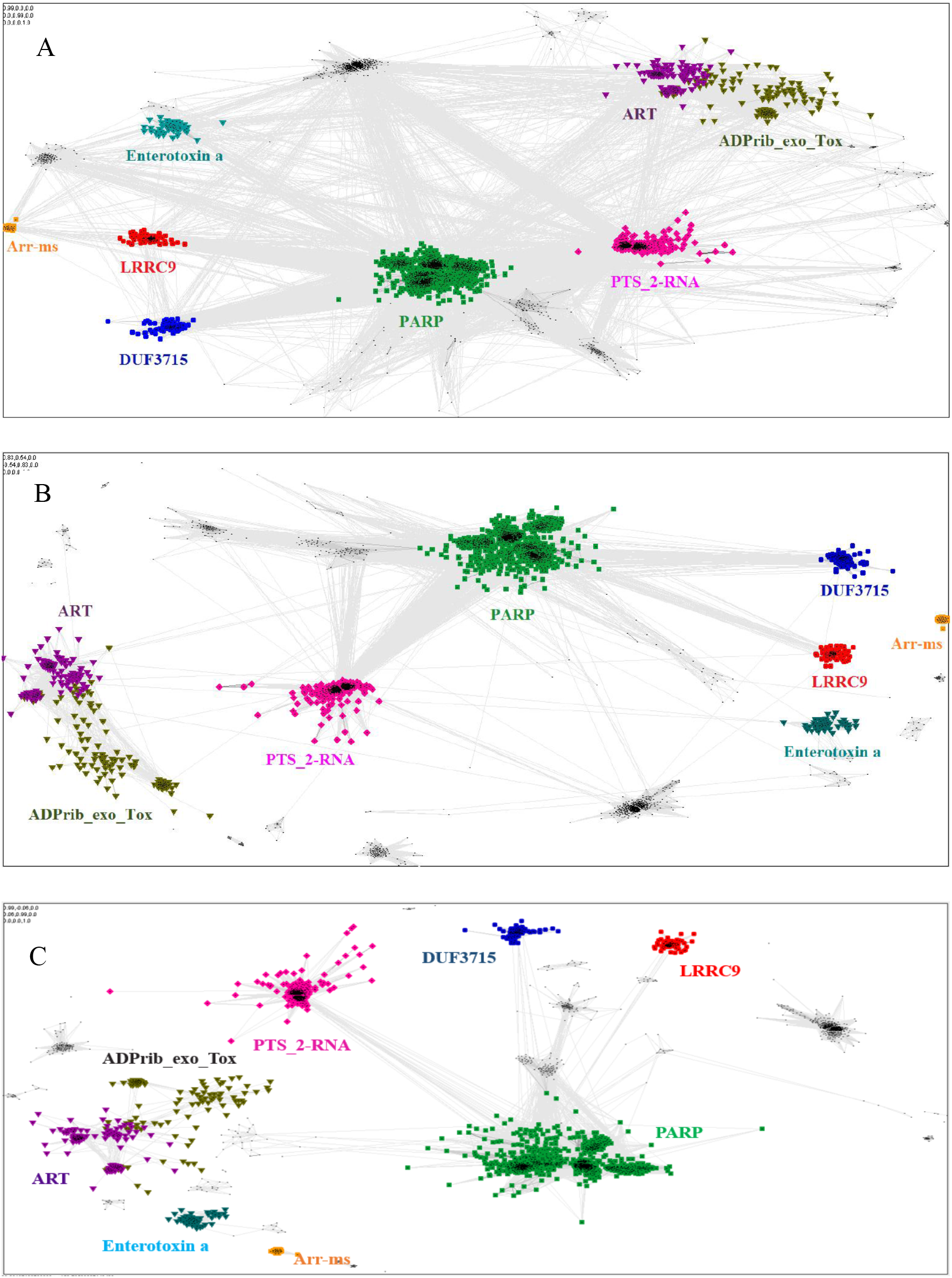
Close and distant sequence similarities between the putative and the known ADP-ribosyltransferase domains, visualized using CLANS algorithm. A. sequence similarities up to BLAST E-value of A: 1, B: at E-value 1e-2, C: at E-value 1e-4

### Putative active site of the novel ART-like family

Catalysis of poly-ADP-ribosylation performed by the members H-Y-[EDQ] clade (including PARP domain family) involves three non-consecutive conserved amino acid residues: His, Tyr and an acidic one (Glu or Asp) that can also be substituted by Gln. For consistency, these motifs will be called motifs I, II and III, respectively. Histidine, occurring in the conserved motif I Hx[ST] [1] within β-strand 1, is responsible for binding the 2-OH group of the adenosine ribose of NAD^+^ and NH_2_ group of the nicotinamide amide *via* hydrogen bonds [3]. Tyrosine (motif II), localized within β-strand 2 [1], interacts with nicotinamide moiety whereas glutamate of β-strand 5 (first position in [QE]x[QED] motif according to Aravind 2018) seems to play a role in maintaining stability of the furanosyl oxocarbenium intermediate [3].

Multiple sequence alignment comparison allowed us to identify few similarities and substantial differences between sequences of PARP catalytic domains and ART-like domains, which we identified in human LRRC9 protein sequence in its central part, from position 392 to 625. The tyrosine residue (motif II), one of the catalytic triad elements in PARP, was found to be replaced by a weakly conserved Glu (E525) among LRRC9-ART domains. Also, His and Ser residues (from the motif I, His-x-Ser, positions 474-476 in the logo in Fig. 1), highly conserved among PARP enzymatic domain members, were not conserved in LRRC9. The His residue was most often substituted by Tyr. Glutamate, the third element of the catalytic triad (motif III, position 604 in the sequence logo), was replaced by a poorly conserved Lys (Fig. 1). In PARP1, PARP2, PARP10, PARP12 and PARP15 sequences, there is also another important Tyr residue (Tyr932 in human PARP10) that is responsible for nicotinamide stacking [82]. Some authors [82] include this second Tyr in the group of conserved residues being the hallmark of PARP ART-domains [82], hence we will use the term catalytic tetrad HYYE, representing the non-contiguous catalytic motif H-x(n1)-Y-x(n2)-Y-x(n3)-E. According to our results, this second tyrosine was conserved in PARP (at the logo position 538, Fig. 1A) and not conserved in the LRRC9 ART-like domain (Fig. 1B). Lack of most of the catalytic amino acid residues, evolutionarily conserved in PARP catalytic domains, suggests that LRRC9 ART-like domains may be pseudoenzymes. Interestingly, the FFAS03 sequence alignment between human LRRC9 ART-like and PARP10 ART domains, especially in its central part is not unequivocal; there are some suboptimal alignment paths to be considered (Suppl. Fig 1). A PARP region flanked by two tyrosines belonging to catalytic tetrad HYYE is aligned to a poorly conserved region 525-538 of the LRRC9 ART-like domain, but three positions upstream and downstream of this fragment there are two distinct motifs of LRRC9-ART domain: PR[IL] and [DE]xxxFRHG, respectively. This suggests that sequence-based alignment may be inaccurate in this region and one of both above-mentioned LRRC9 conserved motifs may in fact be involved in the active site. Also, the strongly conserved D-x(5)-PEY-(2)-EFEY motif in LRRC9 (logo positions 609-623), although not aligned by FFAS to PARP active site, may be speculated to be functionally important.

### Structural analysis of the homology model

To answer the question if the observed distant sequence similarity between LRRC9-ART and PARP families can translate into functional similarity, we focused first on the known structural determinants of ADP-ribosyltransferase activity [83] and relating these to LRRC9-ART features. In classical PARPs, like PARP1 and PARP2, possessing a poly-ADP-ribosylation (PAR) activity, besides the canonical H-x(n1)-Y-x(n2)-Y-x(n3)-[EI] tetrad mentioned above, there are two other key positions associated with structural and functional roles of these proteins: glycine residue with nicotinamide anchoring function and an additional catalytic amino acid, lysine [83]. In PARP10 and PARP12 with mono-ADP-ribosylation (MAR) activity, the glutamic acid is substituted by isoleucine and instead of a catalytic lysine, there is a leucine or tyrosine residue. Investigating profile-profile alignments for human LRRC9 and PARP10 ART domains one can easily notice that only the first of the canonical tetrad residues is conserved, although substituted (H->Y). Histidine and glycine responsible for nicotinamide anchoring and binding are replaced by tyrosine and valine (Figure 5). Interestingly, in LRRC9-ART at position 532 there is a weakly conserved Lys, as in PARP family members with poly-ADP-ribosylation (PAR) activity. However, instead of a second Tyr (logo position 538) and Glu residues (604), there are poorly conserved Glu and Lys. Figure 6 presents a structure model of LRRC9 ART-like domain putative ligand binding site with hypothetical ligand molecule location according to COFACTOR predictions. COFACTOR is a functional annotation tool based on threading of the protein structure query through the BioLiP function database of ligand-protein binding interactions by local and global structure matches to identify functional sites and homologies [84]. COFACTOR suggests the best candidates fitting into the ligand binding site of a given structure. Adenosine monophosphate (AMP) turned out to be among the best scoring ligands for the LRRC9-ART structure model while NAD, a typical ligand for ART domains, was absent from the predicted ligand list. AMP molecule, which differs from NAD by the lack of nicotinamide moiety, is smaller and probably fits better in the LRRC9-ART model binding groove. The docked AMP molecule pose was deduced from homology to crystal structure of the catalytic domain of *Pseudomonas aeruginosa* exotoxin A (PDB ID: 1DMA) which catalyzes the transfer of ADP ribose from nicotinamide adenine dinucleotide (NAD) to elongation factor-2 in eukaryotic cells and hence inhibits protein synthesis. These docking results do not suggest that AMP is a physiological ligand of LRRC9-ART, rather, they provide a likely binding pose of the AMP moiety of NAD.

Replacing the catalytic Glu residue of the HYYE tetrad by Ile is characteristic for PARP family members having mono-ADP-ribosylation (MAR) activity instead of poly-ADP-ribosylation (PAR) [82]. Among FFAS/HHpred hits for LRRC9-ART domain, the only structures that showed up were proteins with MAR activity and PARP13 which has no confirmed ADP-ribosylation activity. In the sequence logos of ART and LRRC9-ART, there are no common conserved motifs between the two domain families. In LRRC9, the second tyrosine of HYYE tetrad is replaced by glutamine 525, which is noticeably less conserved. However, five residues further towards the N-terminus, a conserved motif [ED]-x-P-R-[IL] can be seen. At position 532 in LRRC9-ART domain logo which corresponds to catalytic PARP Lys (substituted by Leu in MAR enzymes) there is a lysine, which could take over some catalytic function in LRRC9 protein. Another highly conserved motif of the PARP family, YxEY[VI][VI][FY], containing catalytic Glu of the HYYE tetrad does not align to a relatively similar motif visible in the LRRC9-ART domain EY[VI][VI]E[FY]EY (logo positions 616-623). As mentioned earlier, these motifs may correspond to each other, and the misalignment may be just an artifact.

Figure 6 shows the active site of a modelled LRRC9-ART structure, with AMP molecule docked in the predicted active site. Fig. 6A presents the same residues as Figure 5 but colored according to conservation in the LRRC9-ART family. Remarkably, only tyrosine 474 and valine 475 (out of the residues aligned to the catalytic tetrad) are quite strongly conserved in the whole family. The rest of LRRC9 amino acids aligned to classical HYYE tetrad residues are not highly conserved. Fig. 6B presents all strongly conserved residues in the neighborhood of the putative LRRC9 ART-like domain active site. It demonstrates that there are some other conserved amino acids within 5 Å of predicted ligand location, like Phe473 and Leu472 (near YV dyad aligned to PARP HG), Ile490, close to adenosine ring of the docked ligand, Phe495 (part of conserved ExGF motif flanking ligand position near N-ribose ring) and Trp605 (next to Lys604 that replaces the catalytic Glu). Ligand pose predicted by COFACTOR enables only two hydrogen bonds between AMP and residues aligned to PARP ART-like domain catalytic tetrad: between AMP phosphate and Val 476 and between N-ribose ring and Glu 525. This method of docking is only preliminary and can give a general idea of ligand placement, but no detailed information on its interactions. The model presented in surface representation with coloring by within-family conservation (Fig. 6C) shows the conserved residues are grouped in the vicinity of the putative ligand binding groove.

The ART-like domain described here is only one region of the whole LRRC9 protein molecule, which is quite large, with clear similarity to leucine-rich repeats containing domains in its N-and C-terminal parts and a very long helical “spine” in the center. We were tempted to construct the whole molecule model, based on homology to LRR-containing templates and using an already modelled ART-like domain structure. The results are shown in Figure 7AB. In the full-length structure model proposed here, a large groove is surrounded by LRR “horseshoes” from three sides and an ART-like domain from the fourth. The size of the groove seems to suggest a possibility of binding a large molecule, e.g. a protein rather than a small cofactor. It is tempting to speculate that LRRC9, binding some “bait”, rearranges its conformation and activates the enzymatic ART-like domain, thereby triggering a cellular signal. However, the full-length model is only an illustration of possible domain arrangements.

### Taxonomic distribution of the LRRC9-like proteins

The LRRC9 protein is widespread in several of the major eukaryotic lineages. It is found in *Alveolata* and *Stramenopiles* (from the TSAR supergroup [85])*, Cryptophyta*, *Haptophyta, Viridiplantae, Fungi and Metazoa (Opisthokonta*) and *Euglenozoa* while absent from several other main lineages, e.g. *Hemimastigophora*, CRuMS, *Metamonada*, *Malawimonadida*, *Ancyromonadida*, *Ancoracysta*, *Picozoa* (Fig. 3). Thus, this protein appears to be an ancestral eukaryotic feature, that was subject to losses in a number of lineages.

ART-like domain is present in all vertebrate classes as well as in 11 non-vertebrate animal phyla (*e.g. Cnidaria, Echinodermata, Brachiopoda, Mollusca, Annelida, Hemichordata, Chordata*). However, notable is its absence in several animal phyla, some of which include important model organisms (e.g. *Arthropoda*, *Nematoda*, *Porifera*; see Fig. 3). Also, strikingly, although LRRC9-ART domains are found in some *Viridiplantae*, e.g. mosses, green algae, liverworts, and club mosses, they are absent from flowering plants with the exception of the water lily *Nymphaea colorata*. Notably, water lilies belong to *Magnoliophyta*, a taxon thought to have diverged earliest from the lineage leading to most extant flowering plants [86]. Altogether, this taxonomic spread suggests an ancient function for LRRC9, common to many diverse eukaryotes.

### Likely biological processes involving LRRC9

Apart from the ART-like domain, LRRC9 proteins comprise a variable number of leucine-rich repeats (LRRs) flanking the ART domain (Fig. 4). In all the homologs analysed, the LRR-ART-LRR domain architecture is conserved in evolution. The N-terminal region possesses 1-6 LRR repeats while the C-terminal region possesses 11-20 such repeats. The N-terminal repeats and C-terminal repeats are most similar to the LRR repeats from proteins such as Leucine Rich Repeat Containing 23 (LRC23) and TLR4 Interactor with leucine-rich Repeats (TRIL), respectively, however, degree of similarity between LRR motifs from different proteins is generally low and makes it difficult to draw specific functional conclusions. The LRR motifs typically have a length of about 20-30 amino acid residues and are rich in leucines or other aliphatic residues (typically seven per motif, localized at positions 2, 5, 7, 12, 16, 21 and 24 within the consensus LRR sequence) [87]. The LRR motifs may form various combinations of two or three secondary structures, usually involving two stretches of a β-strand, e.g. two β-strands and an α-helix. The LRR repeats typically fold into a horseshoe-like conformation composed of helices present on its convex face and parallel β-sheet localized on the concave one [88]. LRR motifs determine functionality of LRR-containing proteins, enabling them to make protein-protein interactions [87] as well as to bind various target structures such as small molecule hormones, lipids and nucleic acids [89]. The non-globular shape of LRR motifs may facilitate the contact between LRR-containing proteins and the target structures, increasing the interaction area [87]. For example, ribonuclease inhibitor is able to bind pancreatic ribonucleases and inhibit their enzymatic activity. This interaction may affect RNA turnover in the context of angiogenesis [90].

**Fig. 4.**
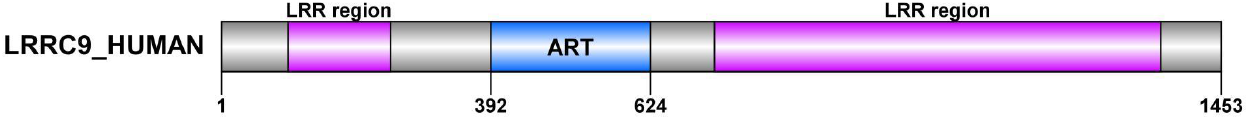
Domain organization of LRRC9 proteins (created using the DOG tool).

**Fig. 5.**
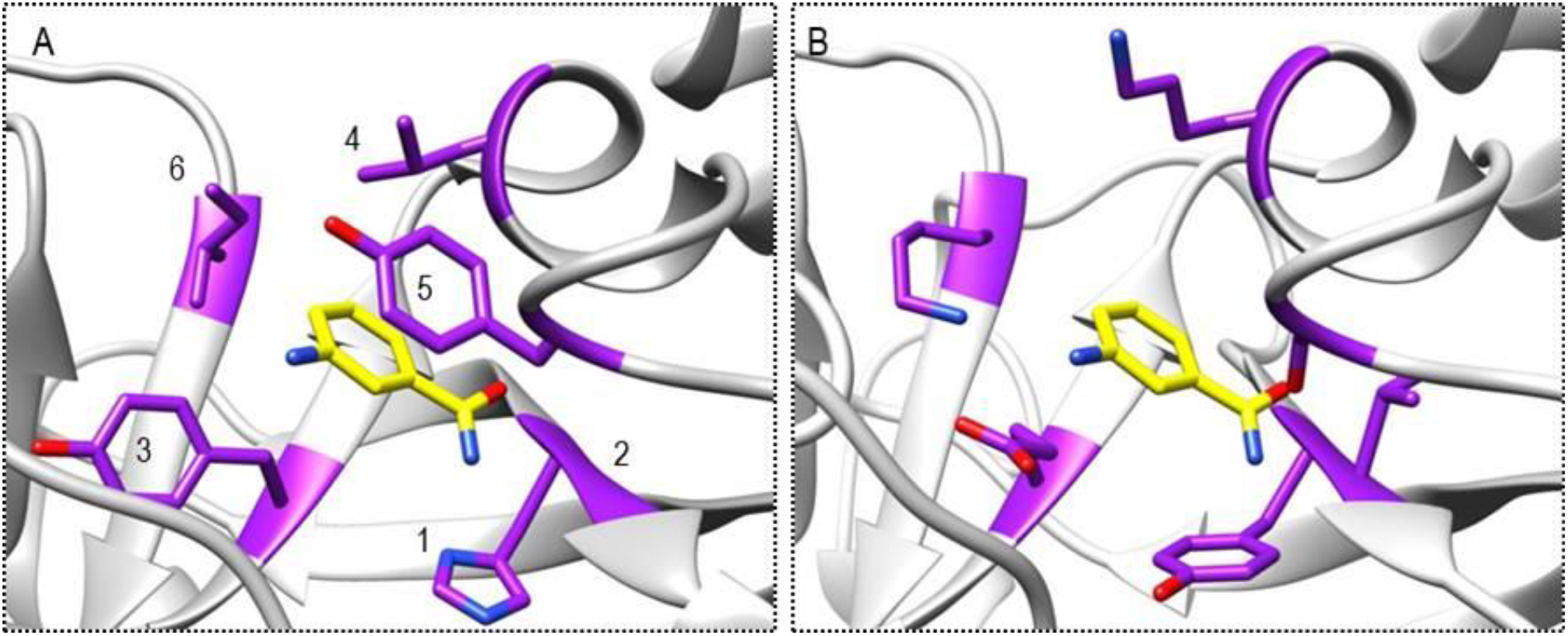
Comparison of active site in the template structure (PARP10, 3HKV) (A) and modeled human LRRC9-ART domain. (B). Canonical PARP tetrad residues (listed in the table below) and their counterparts in the modelled structure are depicted in stick representation. The ligand is 3-aminobenzamide, a PARP inhibitor (yellow).

**Fig. 6.**
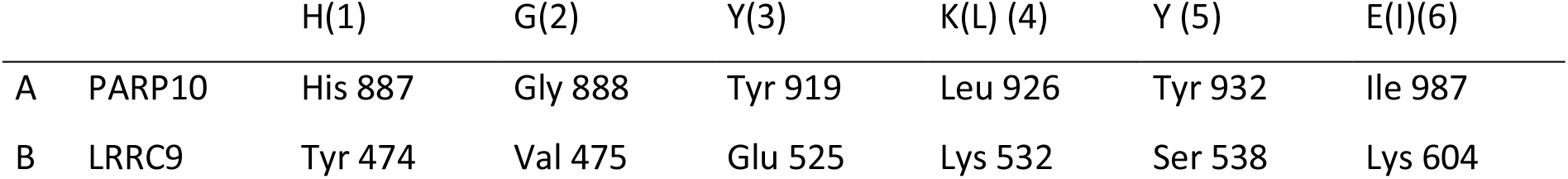
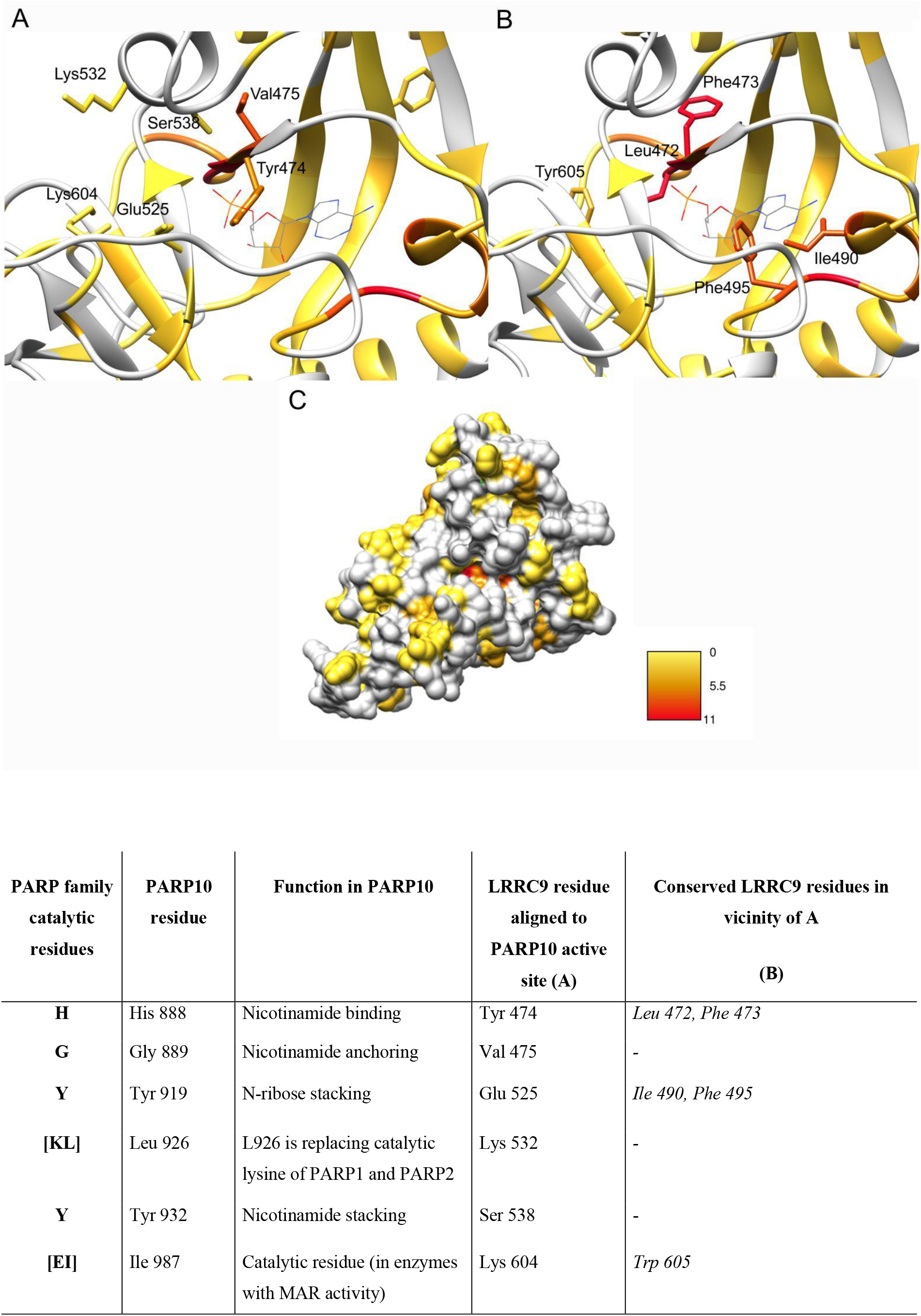
Human LRRC9 homology model in ribbon representation colored according to amino acid conservation (red - highest) in multiple sequence alignment of LRRC9 homologues. An AMP molecule docked in location and pose indicated by COFACTOR algorithm. A. LRRC9 residues aligned with PARP10 active site shown as sticks. B. LRRC9 conserved residues located in the vicinity of the active site shown as sticks. C. Surface representation of LRRC9 structure model with visible conserved residues in the putative binding groove.

**Fig. 7.**
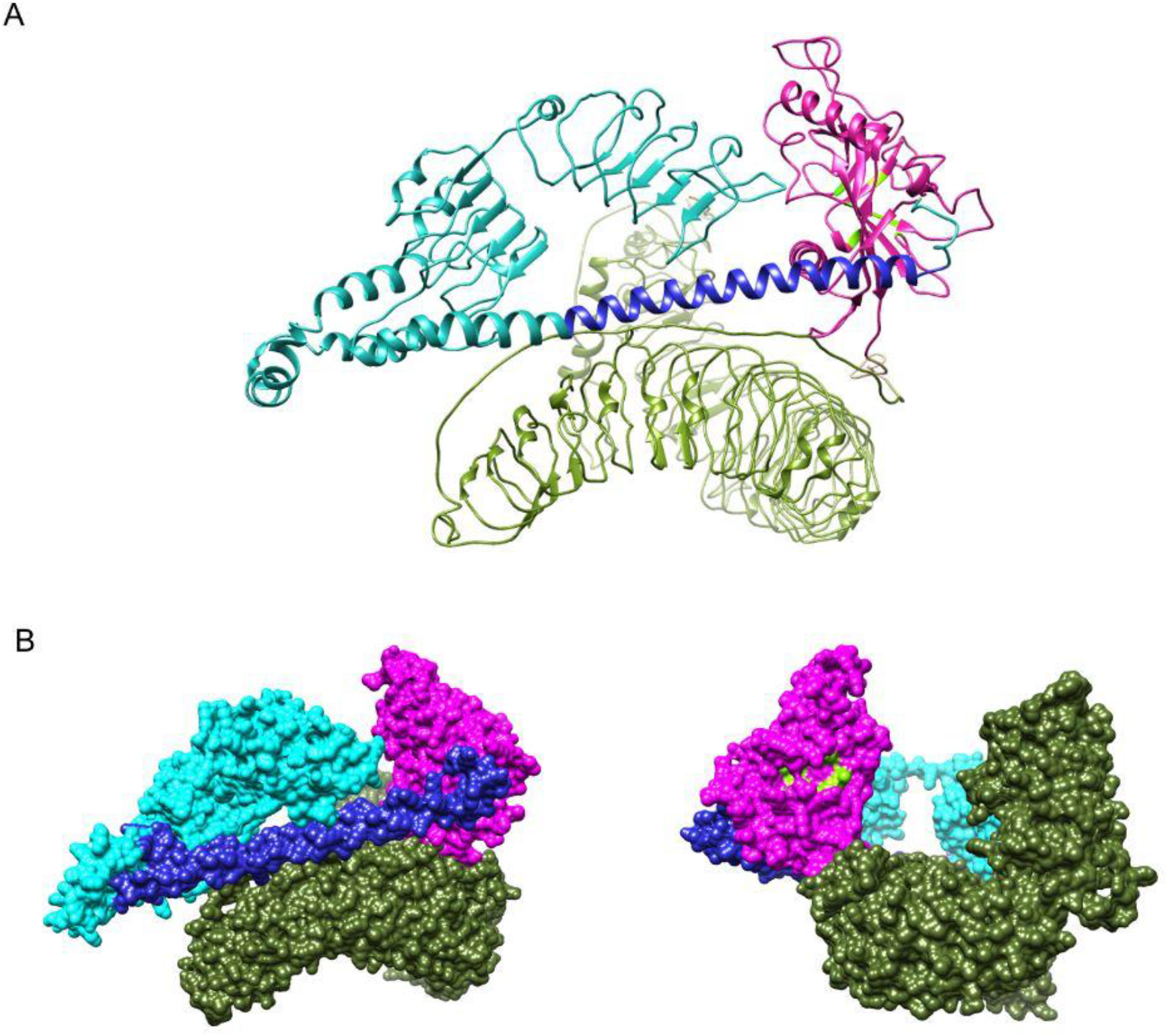
A. Human LRRC9 structure model for the full-length protein in ribbon representation, colored by domain. ART-like domain (magenta) is stacked between two large leucine-rich repeat-containing domains (N-terminal LRR domain with long helical fragment is depicted in cyan and blue, C-terminal LRR domain - in green). B. The same model in surface representation (viewed from two directions).

Employing the LRR motifs, LRR proteins perform various functions. They may play a role in signal transduction and cell development as well as they are able to participate in RNA splicing and effective response to DNA damage [91]. Many LRR-containing proteins are adhesive molecules that perform important functions in processes such as regulation of collagen-fibril formation, osteogenesis, myelination and platelet adhesion at a site of vascular injury. Many proteins with LRR motifs play a role in signal transduction as specific receptors. For example, in response to LPS stimulation, LRR-containing CD14 protein induces tyrosine phosphorylation of intracellular proteins to stimulate antibacterial activity of macrophages. Other receptors exhibit specificity to gonadotrophins such as luteinizing hormone, chorionic gonadotrophin and follicle-stimulating hormone [83].

The LRR domains, which are structures strongly conserved in evolution, occur widely within plant, invertebrate and vertebrate proteins responsible for innate immunity. Due to their ability to mediate protein-protein interactions, LRR repeats determine signaling function of many pattern recognition receptors (PRRs), such as Toll-like receptors (TLRs) and NOD-like receptors (NLRs), by conditioning their affinity to various ligands, i.e. viral, bacterial, fungal and parasite antigens [92, 93]. On the other hand, also ADP-ribosylation is known to be involved in regulation of the host immune response [94, 95]. It may direct intracellular signaling to parthanatos (type of programmed death) [96–98], whereas such scenario implies induction of inflammatory response to infection. Poly-ADP-ribosylation is known to influence the stability of transcripts encoding proinflammatory cytokines [99]. PARP-dependent chromatin modification may counteract progression of viral infection. ADP-ribosylation also influences other innate immune mechanisms such as NF-ĸB expression, phagocytosis and macrophage polarization [100]. Co-occurrence of ART and LRR domains within the LRRC9 protein suggests that they may act together to support a version of host innate immunity. In this context, the LRR domains could perform signal sensors function, and the ART domain might play the effector role.

Overall, in the human proteome there are 234 proteins containing LRR repeats. The molecular functions significantly overrepresented among these proteins include peptide binding (11 proteins, FDR p-value 8E-6), ubiquitin-protein transferase activity (11 proteins), heparin binding (7 proteins) and G-protein-coupled peptide receptor activity (7 proteins). However, as many as 182 human LRR proteins are functionally uncharacterized, according to Panther-db functional annotation analysis. Human proteins containing LRR repeats are involved in processes such as neurogenesis (17 proteins, FDR 1E-5), ubiquitin-dependent protein catabolic process (11 proteins), positive regulation of innate immune response *via* toll-like receptor signaling pathway (10 proteins, FDR 8E-10), cellular response to hormone stimulus (7 proteins) as well as phosphatidylinositol 3-kinase signaling (6 proteins).

Many human LRR-containing proteins undergo extracellular secretion (36 proteins), and many are integral components of plasma membrane (25 proteins).

The ART-like domain combined with LRR motifs has a precedent, since, interestingly, as many as 35 human LRR proteins possess enzymatic domains. Notable here are the NLRP immune sensors/effectors containing NACHT ATP-ase domains, totaling 17 human proteins [101, 102]. Other enzymes with LRR motifs include protein kinases (11 proteins), nucleases and peroxidases. Eight human LRR proteins also possess F-box motifs and are involved in ubiquitin ligase complexes.

Human LRRC9 protein is expressed mainly in the heart, tonsil as well as in the liver (Proteomics-db database). However, no high quality protein expression data is available for LRRC9, hence it is annotated as “existence validated on transcript level” in the NextProt database. This may suggest that LRRC9 protein expression is limited to specific circumstances, and is not easily captured in typical tissue proteomics experiments. LRRC9 mRNA expression is highest within the brain as well as endocrine and male reproductive tissues, i.e. pituitary gland and testis, respectively (Protein Atlas database). According to the DeepLoc prediction, the subcellular localization of LRRC9 is cytoplasmic (likelihood 0.79). Immunohistochemical staining revealed strong cytoplasmic expression of human LRRC9 in the gastrointestinal tract of a few patients suffering from colorectal cancer. Moreover, medium expression of the protein was shown in subjects with liver cancer (expression within liver/gall bladder), stomach cancer, testis cancer, renal and urothelial cancer (kidney/urinary bladder) and melanoma (Protein Atlas database).

For an uncharacterised protein, its physical interactors may shed light on its function. However, according to the BioGRID database, affinity capture-mass spectrometry analysis provided evidence that LRRC9 physically interacts with a single protein, ZFP36L2 [103]. ZFP36L2 is an RNA-binding protein regulating cell cycle [104] that contributes to pathogenesis of pancreatic ductal adenocarcinoma (PDAC). ZFP36L2 is responsible for an increase in cancer cell aggressiveness [105]. A Kaplan-Meier survival analysis (Protein Atlas) shows that LRRC9 mRNA expression is positively correlated with survival among patients with pancreatic cancer (p=0.0034), although the gene is not classified as prognostic. This may suggest an antagonistic relation between LRRC9 and ZFP36L2 activities in the context of ZFP36L2-dependent promotion of PDAC progression. However, the LRRC9-ZFP36L2 interaction was only reported in a high-throughput experiment, and further investigation is required here.

## Conclusions

Pseudoenzymes are characterized by the lack of enzymatic activity despite the common evolutionary origin and structural similarity to catalytically active homologues [106–108]. Pseudoenzymes do occur among PARP family members, including PARP13 that is distinguished by amino acid substitutions and structure changes preventing catalysis. Similarly to LRRC9-ART, PARP13 catalytic domain is characterized by lack of conserved His and Glu residues [83]. It has been shown that substitution of catalytic His with Ala within the ART domain of PARP1 substantially decreases its catalytic activity [109]. The results were consistent with previous studies on diphtheria toxin, another member of H-Y-[EDQ] clade. Replacement of catalytic His reduced the toxin’s activity by at least 70-fold [110]. Thus, the LRRC9-ART family, lacking most of the ART active site, can be hypothesized to be pseudoenzymes. However, one cannot exclude the possibility that this ART-like domain has retained or regained the ART catalytic function, or acquired a novel enzymatic activity, in both cases employing an atypical and/or migrated active site. Such a scenario has been recently reported for apparent pseudokinases, SelO and SidJ, that were shown to be AMPylases and polyglutamylases, respectively [57, 111].

On the other side, in eukaryotic cells, poly-ADP-ribosylating proteins may perform some of their functions without utilizing the enzymatic activity. For example, PARP1 participates in NF-ĸB-dependent gene transcription *via* two different mechanisms but only one requires poly-ADP-ribosylation whereas the second seems not to involve PARP1 enzymatic function and depends on the structure of the protein [21, 112]. Both a pseudo-enzymatic character of LRRC9-ART, and migration of the active site are possible for this intriguing novel family. Experimental studies are needed to confirm one of those hypotheses. The unique, conserved domain architecture of LRRC9, suggests that this mysterious protein family could be involved in a defense mechanism, with some analogies to the innate immune system, coupling within a single molecule the detection of foreign objects (LRRs) and downstream signalling (ART domain).

## Acknowledgements

We thank Drs Vincent Tagliabracci, Miles Black, Anna Muszewska and Marcin Grynberg for critical reading of the manuscript. K.P. was supported by the Polish National Agency for Scientific Exchange scholarship PPN/BEK/2018/1/00431 and by the Polish National Science Centre grant 2019/33/B/NZ2/01409.

